# Investigating carbohydrate-rich feed supplements and the feeding preferences of European starlings (*Sturnus vulgaris*)

**DOI:** 10.1101/2022.02.08.479625

**Authors:** Dylan J. Brown

## Abstract

In the current ecological and ornithological environment in the continental United States, the European starling (*Sturnus vulgaris*) is one of the most common exotic bird species. After being introduced on the East Coast of the United States, the species spread rapidly across regions and ecosystems, significantly affecting the greater ecology of the nation and often disturbing local species populations. To better understand the behavior and preferences of this species, a study was devised to test the field responses of a population of European starlings in Lyon County, Kansas over a total period of 144 hours (during 48 individual observation periods) to assess and yield knowledge pertaining to their behavior and feeding patterns within a suburban environment. For this experiment, four feed alternatives (i.e., one control and three experimental) were prepared to test the impact of the utilization of carbohydrate-rich feed supplements on the degree to which the starlings would frequent a feeding location. The different feed offered to the local starlings would consist of either 300g of conventional bird feed, 200g of conventional bird feed and 100g of the supplement, 100g of conventional bird feed and 200g of the supplement, or 300g of the supplement, with the conventional bird seed acting as the control. The data suggest that the differences observed with relation to the feed alternatives were negligible, as demonstrated by the starlings’ preference for the different supplement concentrations (according to their feeding frequency for the four alternatives).

## 1 Introduction

Throughout many parts of East-Central Kansas, the European starling (*Sturnus vulgaris*) is a common exotic species. Although not originally from the Americas (as their common name suggests), the European starling was introduced to the United States in the late nineteenth century^1^. Over time, the starling has become a common species that is distributed not only across Kansas but also across the North American continent^2^. In this way, understanding their tendencies, preferences, and behaviors is of paramount importance not only to the biological and ornithological communities but also to the fields of conservation and natural study^3^. Moreover, comprehending ways in which the attraction and luring of European starlings can be made more effective can yield important insights that may be useful and applicable with regard to related species and other exotic bird varieties.

At its core, the study aims to better ascertain whether European starlings respond to the introduction of carbohydrate-rich supplements to feeders when compared to more conventional types of bird feed. Likewise, this quantitative study is structured to yield insights that will allow the greater community to better understand how the European starling in the East-Central Kansas region responds to nontraditional forms of birdfeed. In addition to the use of purely food-based preferences, steps were also taken to consider the role of inquisitiveness and curiosity (i.e., core features of starling behavior) during the introduction of the carbohydrate-rich supplements^4,5^. Furthermore, due to the multi-faceted elements that affect ornithological observations that involve the use of birdfeeders in suburban areas, additional considerations and precautions were added to the methodological side of the study in order to ensure that more meaningful conclusions could be drawn from the observations made during the study (such as the standardization of bird feeder size and placement) while prioritizing the health of wild organisms and the greater ecosystem.

With regard to the novelty of the study, no other similar investigation has been undertaken in the existing literature; moreover, the geographic location of the study allows for potentially unique and novel findings with regard to the behavior of European starlings. In a similar vein, the ecological stressors, conditions, opportunities, and limitations faced by starlings in East-Central Kansas may differ from those found in other parts of the nation. For this reason, it is crucial for this study to be undertaken in order to better understand the core behavioral features and preferences of this exotic species, which is often categorized as invasive and destructive to local ecological systems.

In a practical sense, this study can help to advance the collective knowledge in the field of animal behavior (viz., concerning *Sturnus vulgaris*) by helping researchers to understand elements and strategies that may or may not work in the pursuit of attracting starlings to certain locations for population control, as the exotic species is exempted from the Migratory Bird Treaty Act and is scheduled for management and control by civilians without a federal permit^6^. Therefore, the decision to undertake a novel study that investigated and analyzed current conditions and preferences in the behavior of European starlings was unequivocally necessary and is aimed to better understand an exotic bird species that is especially common in the continental United States.

## 2 Prior Considerations & Ethics

Prior to beginning the study in the field, it was important to understand the particular features and idiosyncrasies related to the practical study of European starlings. For example, findings regarding the European starling and its preferences when eating from birdfeeders have shown that the species does not respond well to being fed from tube feeders; however, they do prefer feed compositions that include millet and cracked corn^7,8,9^. For this reason, efforts were taken to eliminate this type of feeder from the methods of the study and use types of food, when possible that would be attractive to them. Furthermore, knowledge about the European starling has yielded information regarding their preferences when they use birdfeeders; specifically, like many other bird species, the European starling prefers foods such as sunflower seeds^10^. For this reason, birdseed that included the aforementioned preferred foods of starlings was chosen and utilized in this study.

When studying bird species and their behaviors via the use of bird-feeding apparatuses, it is important to ensure that the safety and health of the birds being observed are preserved during every step of the process. Accordingly, research was done beforehand in order to ensure that the processes and methods of the study adhered to best practices in the field of birdfeed administration through the use of birdfeeders^11,12^. Likewise, the safety of the location in which the birdfeeder was used were inspected and verified according to existing literature and recommendations prior to the beginning of the practical steps of the study.

Furthermore, all steps and elements of the study were conducted according to extant laws and procedures and were explicitly structured to abide by the Animal Welfare Act of 1966 and K. S. A. 21-6411 (i.e., Kansas state law). Moreover, all possible assurances and precautions were undertaken to comply with local and state laws pertaining to studies that focus on vertebrate behavior.

## 3 Methods

As a part of the study, an open-style birdfeeder was placed 1.5 m above the ground in a semi-residential area within a suburban community in East-Central Kansas. The same open-style birdfeeder was employed for two distinct groups, namely, one *control group* and three *experimental groups*. The birdfeeders were utilized during different 48-hour periods in order to avoid competition between the two feeding alternatives, which would likely have skewed the data and may have provided a less clear understanding of starling feeding preferences. For the control group, 300g ± 5g of standard (mostly seed-based) birdseed was evenly spread out across the circular, open-style birdfeeder and left in a clearing with moderate bird traffic for 48 hours (per group being observed). A similar process was done with the experimental groups; however, instead of 300g of only standard birdseed, a carbohydrate-rich supplement was added so that the total amount of food still totaled 300g, creating a 2:1, 1:2, and 0:3 birdseed to carbohydrate-rich supplement ratio (i.e., totally substituting regular bird feed by the third experimental group while the control group was entirely regular bird seed).

During the 48 hours during which each group’s birdfeeder was available for the starlings, the birdfeeders were watched and monitored for starling activity. The majority of the monitoring took place over the course of 30-minute intervals during the hours of peak starling activity (i.e., observed to be during twilight hours and during the mornings/evenings). During these monitoring periods, all bird species seen that were not starlings were not counted or included in the final data. At the beginning of each observation period, the feeder was filled, if necessary. Only starlings that came to or circled around the birdfeeder were counted. In this way, the experiment was structured to record only the number of European starlings observed during the monitoring periods (i.e., over the course of the 48 hours when the birdfeeders were available).

## 4 Results

The methodologies were applied during the winter season in central Lyon County, Kansas. The observations made were gained by adhering to the aforementioned methodological structure and the processes for completing the study. After conducting the observations, the data gleaned was compiled into a table:

To help better visualize the data, charts are provided below.

*N*.*B*., Figures 1 & 2 do not plot dependent against independent variables; they are presented in order to provide a better sense of the chronological order and progression of the observation groups among the three experimental and control groups.

**Figure 1.**
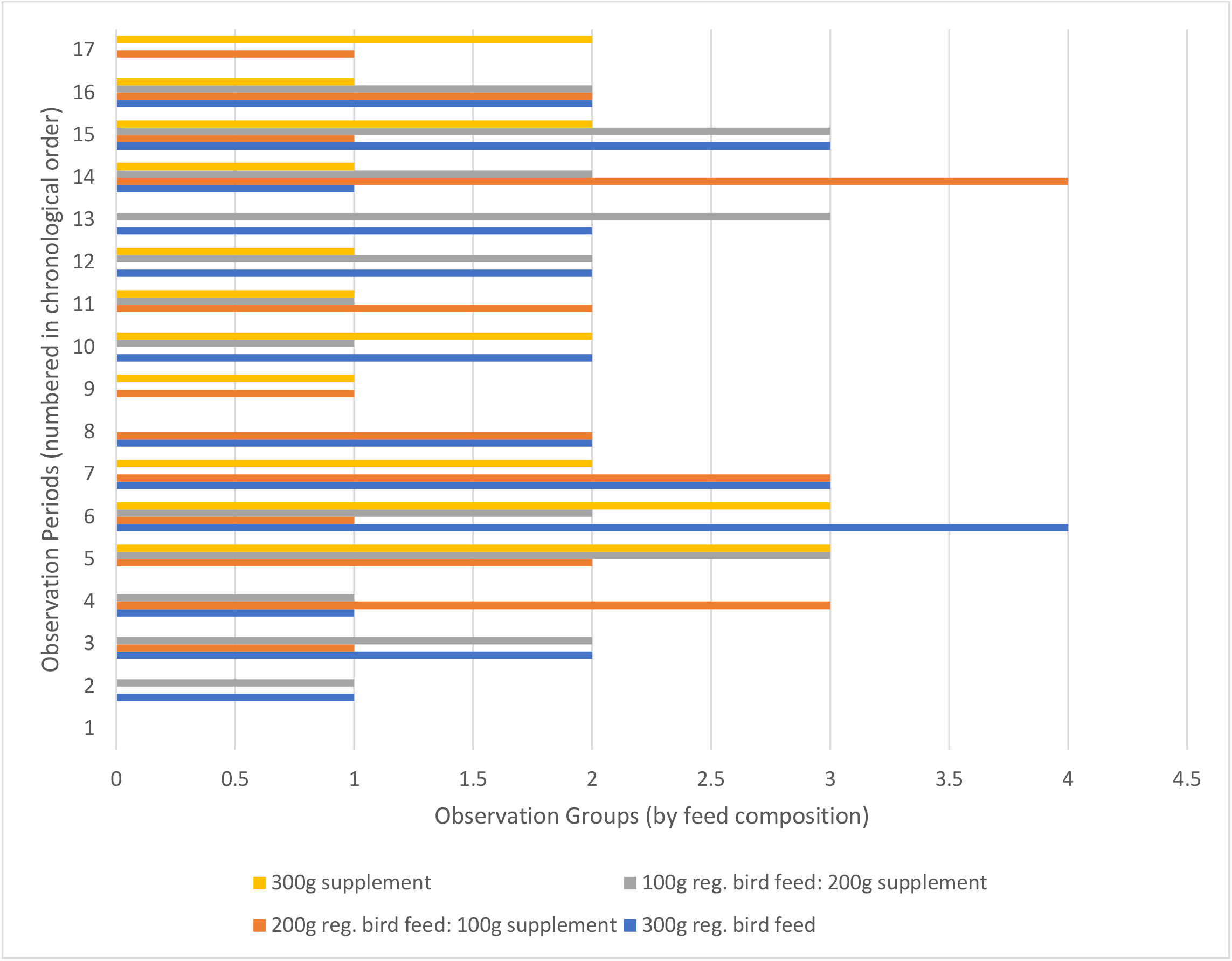
Observation groups (by feed composition) plotted against observation periods plotted against o

**Figure 2.**
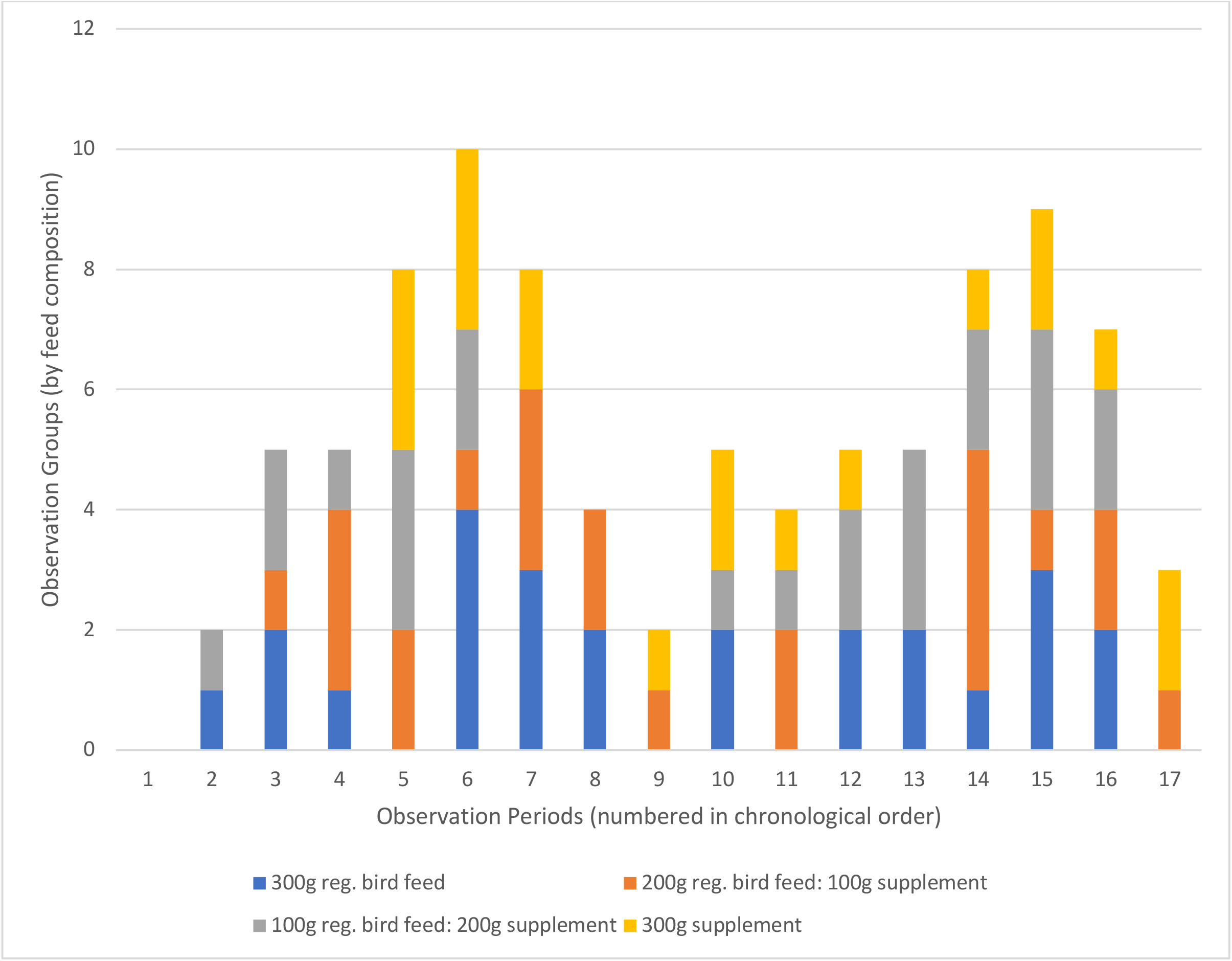
Observation periods plotted against observation groups (by feed composition)

**Figure 3.**
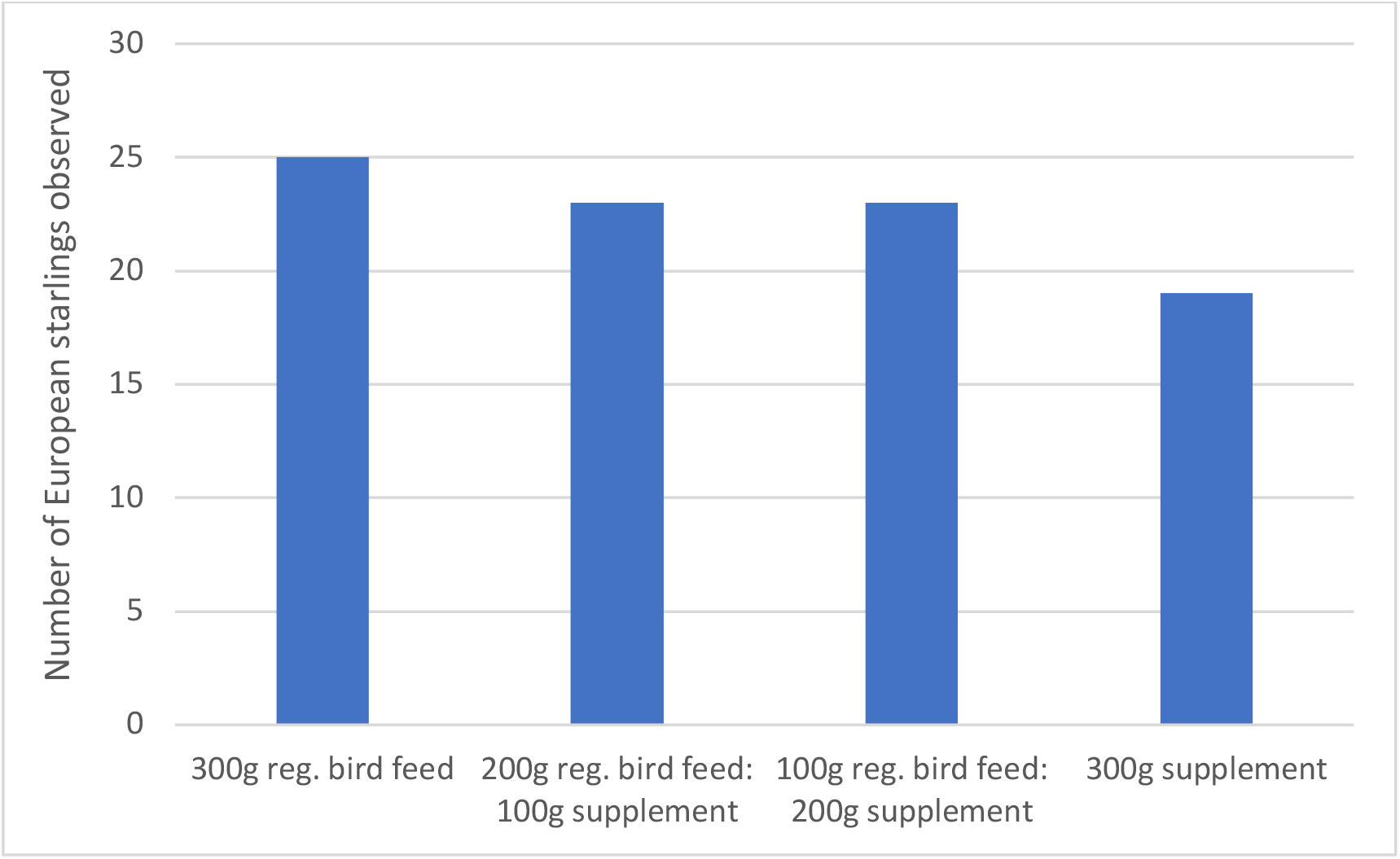
Number of European starlings observed for each feed/supplement group

**Figure 4.**
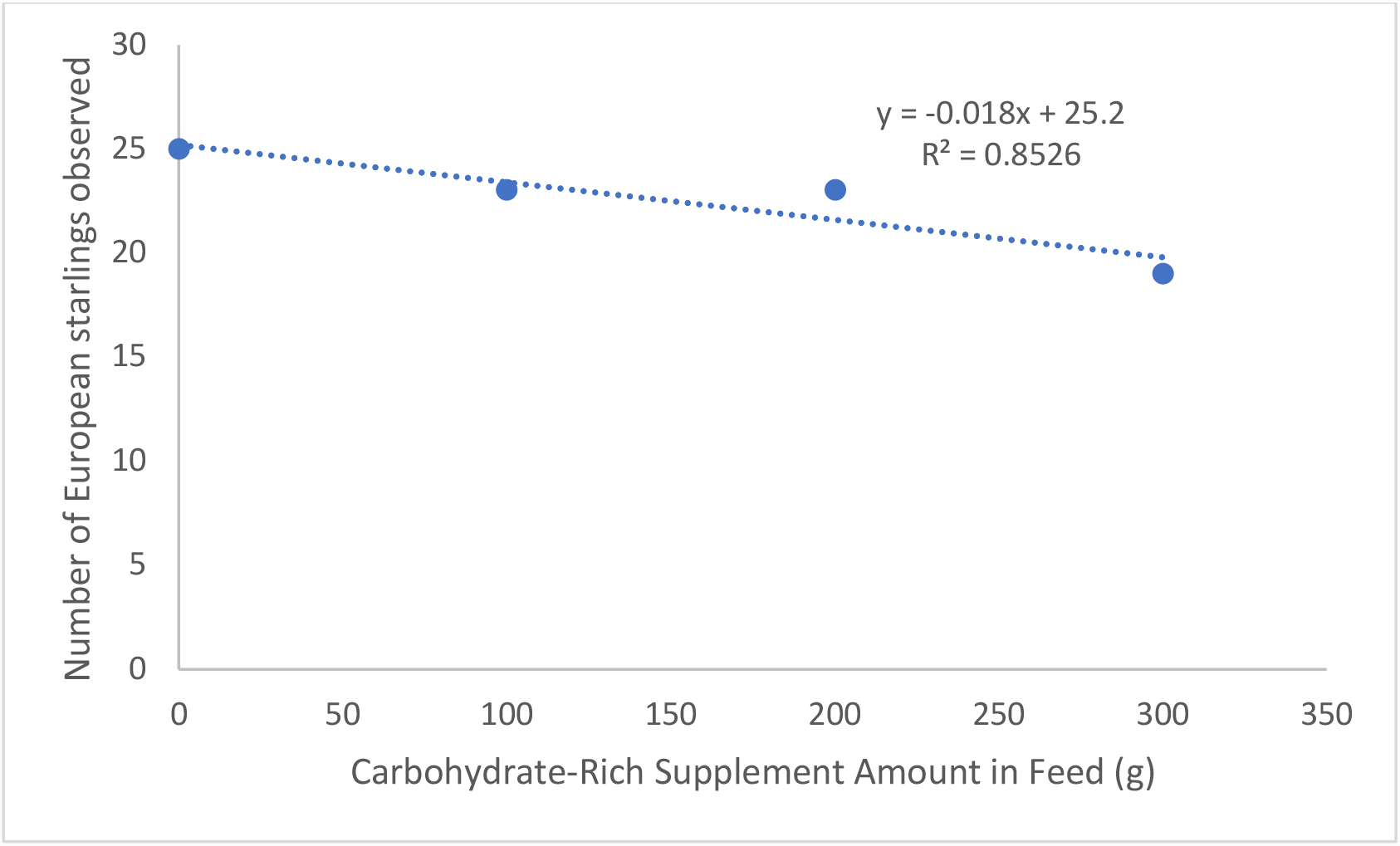
Number of European starlings observed (cumulative) and the amount of carbohydrate-rich supplement in feed alternative (g) with associated trend line and regression statistics

The data presented shows a relatively dispersed data set across the different observation periods and observation groups. Although some cursory assumptions may be drawn, based on the charts above, is difficult to say conclusively how European starlings react to the different feed compositions within the greater context of the study.

Thus, in order to better demonstrate the relationship between the independent variable and dependent variable in this experiment, another chart is provided below that both shows the number of European starlings observed for each group.

To accompany these regression statistics, an ANOVA was also calculated; it is presented in Tables 2 and 3:

**Table 1.** Observations of the number of *Sturnus vulgaris* appreciated over the observation periods per observation group

| Observation Groups | *Sturnus vulgaris* observed |  |  |  |
| --- | --- | --- | --- | --- |
|  | Control Group: Without Carb Supplement | Experimental Group 1: With 100g Carb Supplement | Experimental Group 2: With 200g Carb Supplement | Experimental Group 3: With 300g Carb Supplement |
| Feed Composition | 300g reg. bird feed | 200g reg. bird feed: 100g supplement | 100g reg. bird feed: 200g supplement | 300g supplement |
| Observation Periods |  |  |  |  |
| *Period 1* | 1 | 0 | 1 | 0 |
| *Period 2* | 2 | 1 | 2 | 0 |
| *Period 3* | 1 | 3 | 1 | 0 |
| *Period 4* | 0 | 2 | 3 | 3 |
| *Period 5* | 4 | 1 | 2 | 3 |
| *Period 6* | 3 | 3 | 0 | 2 |
| *Period 7* | 2 | 2 | 0 | 0 |
| *Period 8* | 0 | 1 | 0 | 1 |
| *Period 9* | 2 | 0 | 1 | 2 |
| *Period 10* | 0 | 2 | 1 | 1 |
| *Period 11* | 2 | 0 | 2 | 1 |
| *Period 12* | 2 | 0 | 3 | 0 |
| *Period 13* | 1 | 4 | 2 | 1 |
| *Period 14* | 3 | 1 | 3 | 2 |
| *Period 15* | 2 | 2 | 2 | 1 |
| *Period 16* | 0 | 1 | 0 | 2 |
| Total | 25 | 23 | 23 | 19 |

**Table 2.**
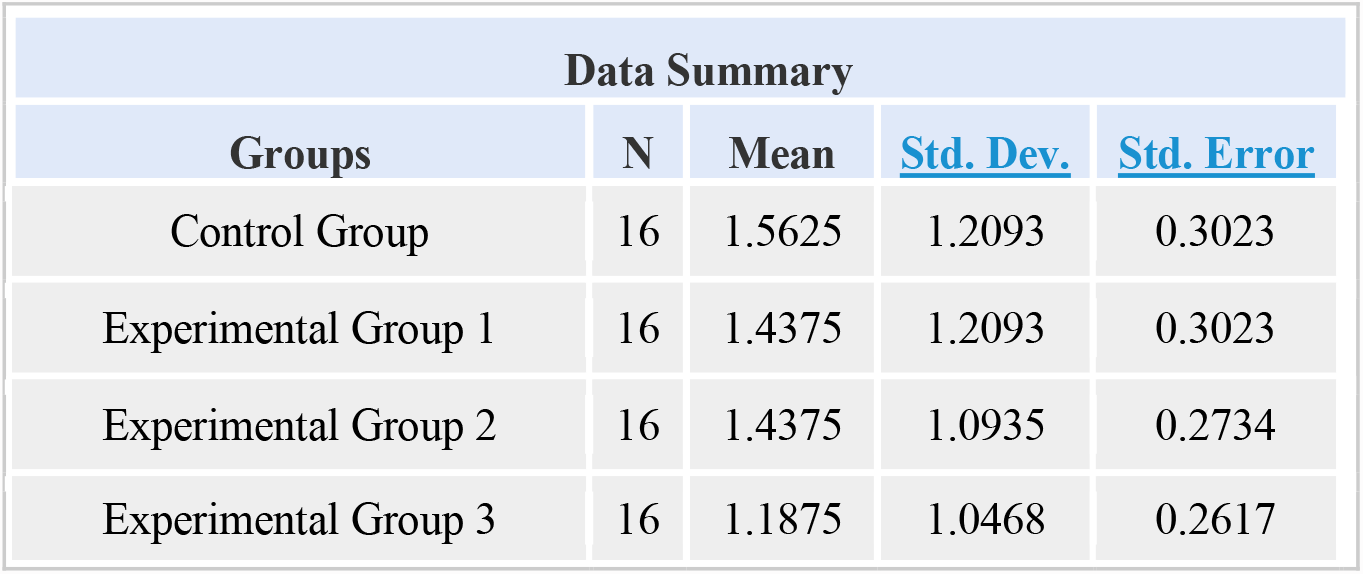
Data Summary for Statistical Calculations for Control and Experiment Groups

**Table 3.** ANOVA Statistical Summary for Degrees of Freedom, Sum of Squares, Mean Square, F-state and P-value

| ANOVA Summary |  |  |  |  |  |
| --- | --- | --- | --- | --- | --- |
| Source | Degrees of Freedom<br>DF | Sum of Squares<br>SS | Mean Square<br>MS | F-Stat | P-Value |
| Between Groups | 3 | 1.1875 | 0.3958 | 0.3035 | 0.8227 |
| Within Groups | 60 | 78.2452 | 1.3041 |  |  |
| Total: | 63 | 79.4327 |  |  |  |

This element of the statistical calculations for the data in this study was central to the quantification and understanding of the core statistical figures of the data set. Notably, it helped to elucidate the average (i.e., arithmetic mean), standard deviation, and standard error within the four distinct groups in the data set.

The use of ANOVA was a crucial step in quantifying and ascertaining the statistical meaning and significance of the data collected. As derived from the calculations, the F-statistic value = 0.30353 and P-value = 0.82272.

## 5 Conclusions

As seen via the presentations of the data and the associated trend line and ANOVA analysis included, there was a general negative slope in the relationship between the feed composition (in terms of the amount of the carbohydrate-rich supplement included in the feed) and the number of starlings observed, the results were determined to not be statistically significant (as seen in the ANOVA P-value calculated). Hence, it is clear the collection of a greater amount of data is necessary in order to draw statistically significant conclusions as to the nature of starling feed preferences and to determine whether any deeper or more significant relationships between these variables exist. Thus, future studies are recommended in order to better ascertain the behavioral feeding preferences of the European starling.

Although significant study results were not able to be gleaned from the data set associated with this study, further research should seek to include larger geographical and population data statistics to better identify any underlying relationships between the variables included in this study. Additionally, future researchers would be wise to include a greater number of potential determiners within the framework of their studies in order to better identify statistically significant relationships that may not have been within the exclusive domain of this study.

Although this research effort did not unequivocally demonstrate a significant relationship between the two variables, it provides a meaningful step forward in helping the larger starling-studying community to gain a sense of the degree to which a significant relationship *does not* exist between this carbohydrate-rich feed composition and European starling feeding preference.

## Acknowledgements

A special word of gratitude is extended to those that helped and supported this study of local Kansas birds; I am deeply thankful for those individuals and mentors who offered both advice, wisdom, and patient understanding during this project, which extended from 2021 through early 2022.

## Author Contribution

The study’s concept was outlined, designed, and tested by Dylan J. Brown. Analysis and critical assessment were also completed by Dylan J. Brown as a part of the project. This manuscript was written by Dylan J. Brown.

